# The structural basis of the oncogenic mutant K-Ras4B homodimers

**DOI:** 10.1101/2020.09.07.285783

**Authors:** Kayra Kosoglu, Meltem Eda Omur, Hyunbum Jang, Ruth Nussinov, Ozlem Keskin, Attila Gursoy

**Affiliations:** Department of Molecular Biology and Genetics, Koc University, Rumelifeneri Yolu, 34450 Sariyer Istanbul, Turkey; Computational Structural Biology Section, Frederick National Laboratory for Cancer Research, National Cancer Institute at Frederick, Frederick, MD 21702, U.S.A; Department of Human Molecular Genetics and Biochemistry, Sackler School of Medicine, Tel Aviv University, Tel Aviv 69978, Israel; Department of Chemical and Biological Engineering, Koc University, Rumelifeneri Yolu, 34450 Sariyer Istanbul, Turkey; Department of Computer Engineering, Koc University, Rumelifeneri Yolu, 34450 Sariyer Istanbul, Turkey

## Abstract

Ras proteins activate their effectors through physical interactions in response to the various extracellular stimuli at the plasma membrane. Oncogenic Ras forms dimer and nanoclusters at the plasma membrane, boosting the downstream MAPK signal. It was reported that K-Ras4B can dimerize through two major interfaces: (i) the effector lobe interface, mapped to Switch I and effector binding regions; (ii) the allosteric lobe interface involving α3 and α4 helices. Recent experiments showed that constitutively active, oncogenic mutant K-Ras4B^G12D^ dimers are enriched in the plasma membrane. Here, we perform molecular dynamics simulations of K-Ras4B^G12D^ homodimers aiming to quantify the two major interfaces in atomic level. To examine the effect of mutations on dimerization, two double mutations, K101D/R102E on the allosteric lobe and R41E/K42D on the effector lobe interfaces were added to the K-Ras4B^G12D^ dimer simulations. We observed that the effector lobe K-Ras4B^G12D^ dimer is stable, while the allosteric lobe dimer alters its helical interface during the simulations, presenting multiple conformations. The K101D/R102E mutations slightly weakens the allosteric lobe interface. However, the R41E/K42D mutations disrupt the effector lobe interface. Using the homo-oligomers prediction server, we obtained trimeric, tetrameric, and pentameric complexes with the allosteric lobe K-Ras4B^G12D^ dimers. However, the allosteric lobe dimer with the K101D/R102E mutations is not capable of generating multiple higher order structures. Our detailed interface analysis may help to develop inhibitor design targeting functional Ras dimerization and high order oligomerization at the membrane signaling platform.

## Introduction

Ras proteins are small GTPases which couple cell-surface receptors to downstream effectors regulating various cellular processes including cell cycle progression, cell differentiation and survival, cytoskeletal organization, cell polarity and movement, and vesicular and nuclear transport. Ras proteins cycle between two conformations: inactive GDP-bound and active GTP-bound forms [1, 2]. The extracellular stimuli lead to the activation of a regulatory protein, guanine nucleotide exchange factor (GEF). GEFs induce the release of guanosine diphosphate (GDP) from Ras and permits binding of guanosine triphosphate (GTP) [3, 4]. Upon GTP binding, a conformational change occurs in downstream effector binding region which allows the interaction of Ras with its effector proteins including Raf kinase, phosphatidylinositol 3-kinase (PI3K), and Ral guanine nucleotide dissociation stimulator (RalGDS) [5–8]. The activity of Ras is downregulated by GTPase activating proteins (GAPs). Ras proteins have intrinsic GTPase activity, which means that they can hydrolyze GTP to GDP. This hydrolysis reaction is extremely slow [2]. GAPs induce the GTPase activity of Ras, thereby accelerates the process. Ras mutations that impair GTPase activity are insensitive to GAPs, rendering mutant Ras proteins persistent in their active GTP-bound state, thereby prolonging downstream signaling associated with oncogenic cell growth [3, 4].

The three human Ras genes encode highly similar proteins: H-Ras, N-Ras and K-Ras. The two K-Ras proteins arise from alternative splicing at their C-termini: K-Ras4A and K-Ras4B [9]. All have 189 amino acids except K-Ras4B that has 188 amino acids. The catalytic domain contains functional P-loop (residues 10-17), Switch I (residues 30-38), and Switch II (residues 60-76) regions, which are responsible for GTP hydrolysis and effector binding. Upon dissociation of GDP and subsequent GTP binding, the conformational change is observed in two regions of Ras; Switch I and Switch II. Switch I is responsible for interaction with GAP and other effector proteins, while Switch II is involved in GEF binding. While the catalytic domain (residues 1-166) of the four isoforms have high identity among each other (~ 89%), the hypervariable region (HVR) of the four isoforms has low sequence identity (~ 8%) (**Fig 1**) [10]. Despite being highly homologous, these isoforms may prefer different binding partners, and have unique physiological functions [11]. K-Ras4B is confirmed as the most frequently mutated isoform in *RAS*-driven cancers (86%), while N-Ras (11%) and H-Ras (3%) are accordingly less mutated isoforms from The Catalog of Somatic Mutations in Cancer (COSMIC) [12]. 98% of oncogenic Ras have amino acid mutations at the active site residues G12, G13, and Q61 [1, 11, 13]. The mutation frequencies vary in K-Ras4B. G12 is the most frequently mutated residue (89%), followed by G13 (9%) and Q61 (1%) residues [14]. G12 most frequently mutates to aspartic acid, G12D (36%), among three frequent mutations G12C (14%) and G12V (23%) [9]. These mutations at conserved sites impair the intrinsic and GAP catalyzed hydrolysis of GTP [15, 16].

**Fig 1.**
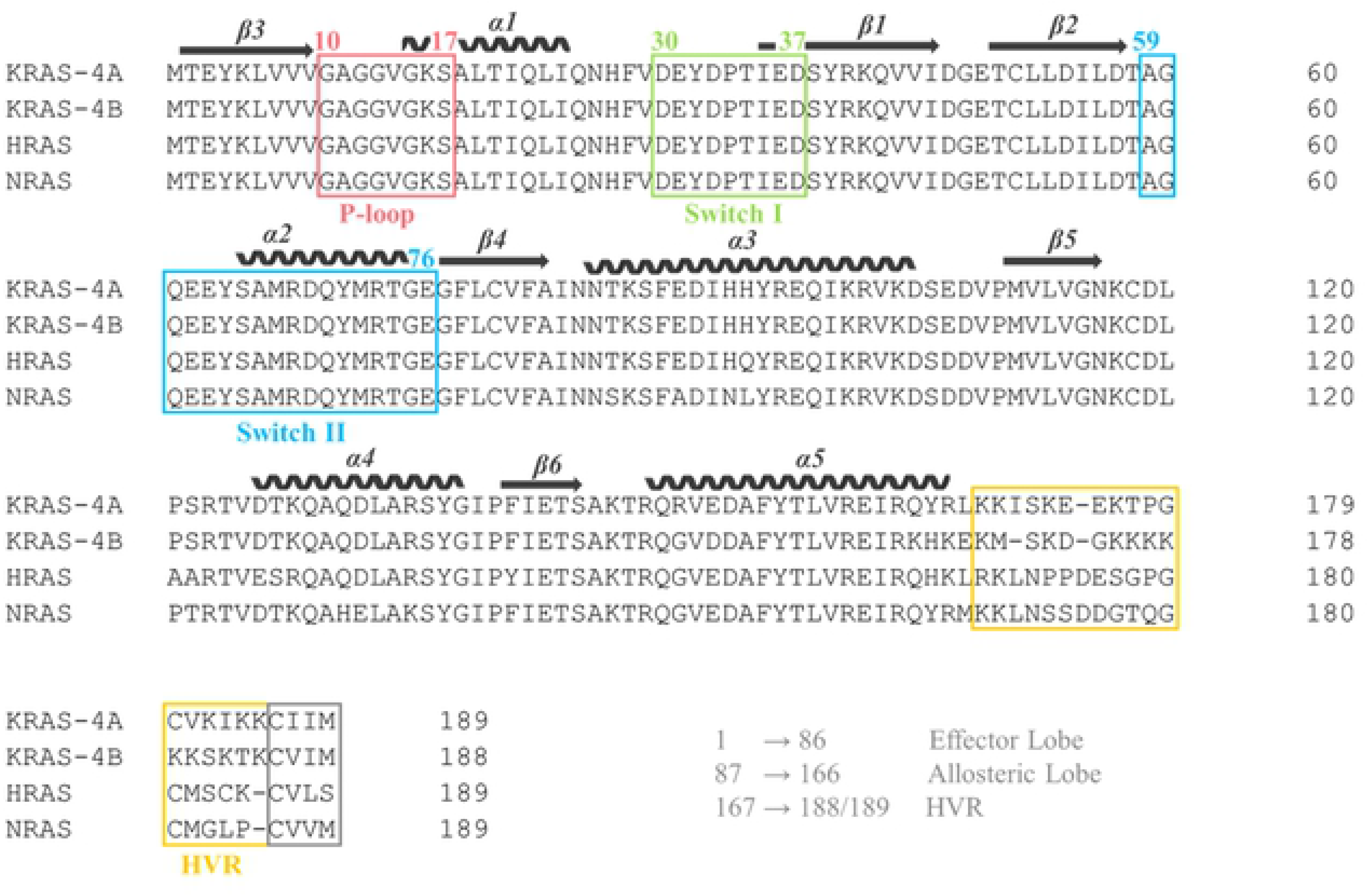
Comparison between Ras isoforms. Sequence similarity of the Ras catalytic domain via multiple sequence alignment of four Ras isoform proteins.

Ras proteins have been defined as monomeric GTPases for a long time. Several studies have provided compelling evidences for existence of their higher order structures [17–25]. Nanoclusters of receptors in cell membranes have been known for a while. N-Ras-GDP was found to form dimers in a model membrane [26]. H-Ras could dimerize on membrane surfaces, and the Switch II region was involved in the dimerization [27]. The Raf kinases are important molecules in Ras signaling pathway. It is known that Raf dimerization plays a critical role in Ras dependent Raf activation [17, 28, 29]. Raf proteins are recruited to the cell membrane upon Ras activation [22, 29–31]. Accordingly, the recruitment results in dephosphorylation of inhibitory sites and the phosphorylation of activating sites within kinase domain. It is believed that Ras dimerization contributes to Raf dimerization and Raf activation [29, 32]. Spencer-Smith *et al.* [33] showed that a synthetic monoclonal protein, binding to the α4-β6-α5 region of H-Ras and K-Ras disrupting Ras dimerization. Ambrogio *et al*. showed that dimerization was required to maintain the oncogenic function of mutant K-Ras [34]. We recently showed that K-Ras4B can form stable dimers through allosteric lobe and this dimerization enhances but not necessary for downstream signaling [19–22]. Despite all these efforts, the mechanism how K-Ras4B dimerizes and promotes Raf activation is yet to be discovered.

We have previously showed that wild-type K-Ras4B can form homodimers through both allosteric and effector lobe dimer interfaces *in silico* and *in vitro* [19–22]. The allosteric lobe dimer interface involves α3 and α4 helices (hereafter referred to as α-homodimer), while the effector lobe dimer interface contains a shifted β-sheet extension between β2 strands (hereafter referred to as β-homodimer) (**Fig 2**). In this study, we adopted both homodimer interfaces from wild-type K-Ras4B dimers and introduced the oncogenic G12D mutation to K-Ras4B dimers. Explicit molecular dynamics (MD) simulations were performed on oncogenic mutant K-Ras4B^G12D^ α-homodimer containing the α3 and α4 helical interface and β-homodimer assembled through a shifted β-sheet extension. To test stability of the oncogenic dimers, we applied two double mutations, K101D/R102E on the α-homodimer interface and R41E/K42D on the β-homodimer interface studied in previous experiments [20], to the oncogenic dimer model systems. Two additional mutant systems, K-Ras4B^G12D/K101D/R102E^ α-homodimer (hereafter referred to as mutant α-homodimer) and K-Ras4B^G12D/R41E/K42D^ (hereafter referred to as mutant β-homodimer) were also subject to explicit MD simulations in solution. Presumably, we expect that the charge converted mutations at the dimeric interface directly interfere with the dimer association. In our simulations, we observed that both oncogenic K-Ras4B^G12D^ α- and β-homodimers are stable, with the oncogenic β-homodimer being more stable than the oncogenic α-homodimer, consistent with our previous study of the wild-type K-Ras4B dimer systems [19, 21]. However, the double mutations R41E/K42D introduced in the effector lobe interface are more disruptive than the double mutations K101D/R102E in the allosteric lobe interface. Both mutant α- and β-homodimers are less stable than the oncogenic homodimers, being prone to interrupt dimer association.

**Fig 2.**
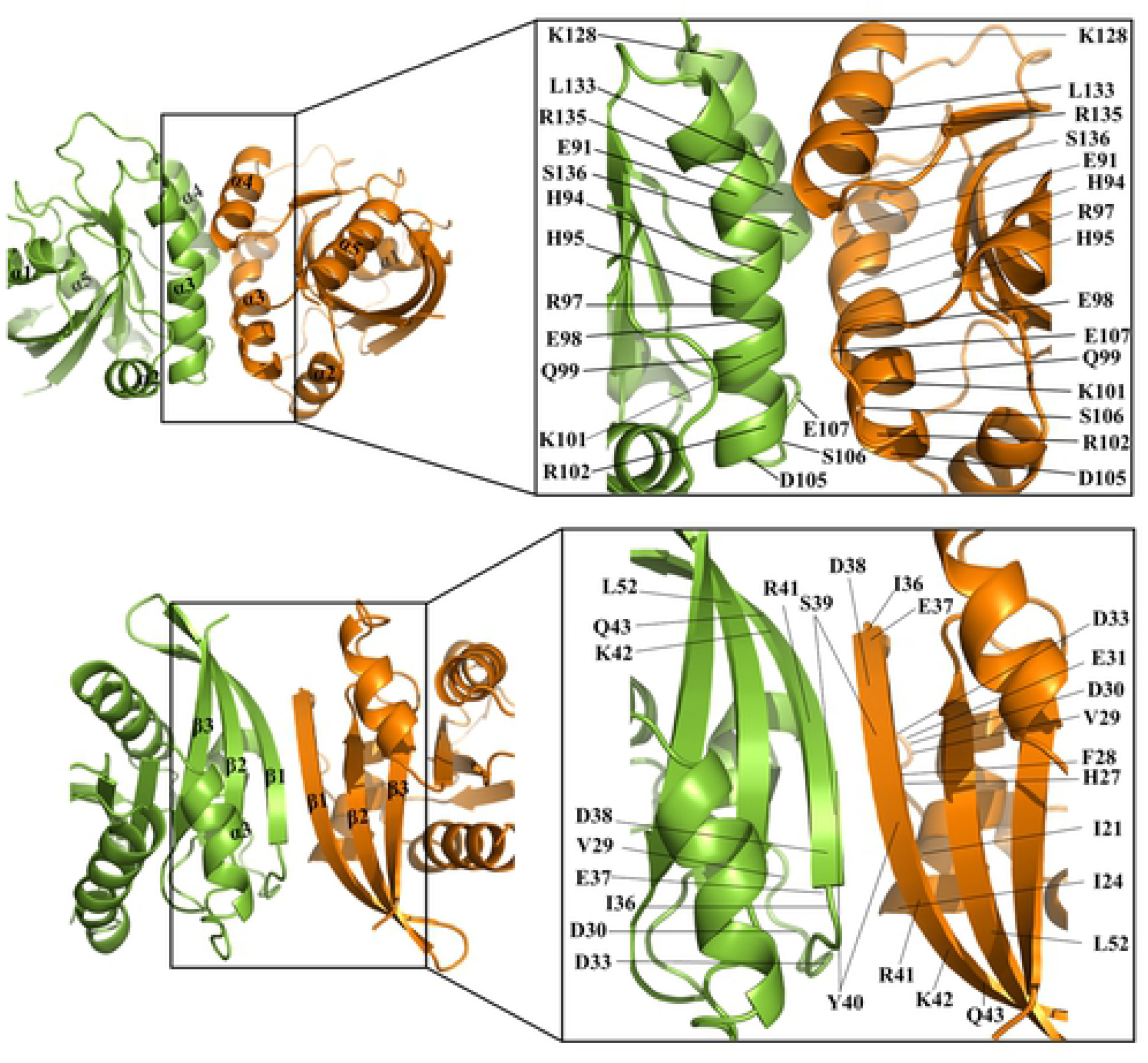
Structure of K-Ras4B homodimers and interface residues. The K-Ras4B α-homodimer involving the symmetric α3-α4/α3-α4 helical alignment (upper row) and the β-homodimer containing a shifted β-sheet extension between β2 strands (lower row).

## Materials and methods

### Computational prediction of K-Ras4B dimers

The dimeric structures of the K-Ras4B were modeled by PRISM [38–40], which is a template-based protein-protein structure prediction algorithm. The outputs of PRISM were ranked based on the binding energy scores (BES). The GTP-bound K-Ras4B structure was obtained from the Protein Data Bank (PDB) (PDB ID: 3GFT). Then, the interface regions of the predicted dimers were identified by HotRegion which also gives the predicted hot spot clusters [41]. HotRegion identifies the important regions for the stability of protein-protein complexes.

### Determination of the residues to be mutated

The atomic interactions of K-Ras4B homodimers in the GTP-bound state were investigated to the interface residues to be mutated. The change in binding free energy (ΔΔ*G*) upon mutations was calculated by FoldX which estimates the stability effect of a mutation by using an empirical method [42]. If the energy value of a mutation is ΔΔ*G* > 0 kcal/mol, then that mutation will destabilize the structure, if the reverse effect is obtained then that mutation will stabilize the structure.

### Atomistic MD simulations

A total of four initial configurations, two α-homodimers and two β-homodimers of K-Ras4B-GTP, were subjected to the MD simulations. Our simulations closely followed the protocol reported in previous studies [21, 22, 43–45]. All-atom additive CHARMM36 force field [46] was used, and simulations were performed by NAMD [47]. Each system was run for 300 ns resulting in a total of 1.2 μs MD simulations. K-Ras4B control and mutant systems were neutralized by addition of 56 Na^+^ and 40 Cl^−^, 64 Na^+^ and 40 Cl^−^ ions, respectively. Mg^2+^ ions are kept. Our protein complexes were simulated in 90 × 90 × 90 Å^3^ virtual water boxes created by using TIP3P explicit solvent model [48]. Before production runs, 10,000 steps of minimization and 50,000 steps of dynamics runs were applied to our system. Dynamics were run under NPT ensemble. The step size was 2 fs. To calculate the long-range electrostatic interaction, particle mesh Ewald (PME) method was used. In the production runs, the Langevin temperature control maintained the constant temperature at 310 K, and the pressure was kept at 1 atm. The simulated trajectories were analyzed using CHARMM [49] and Chimera [50].

### Binding free energy calculation for the Ras dimers

To investigate the strengths of the interactions within the systems, we calculated binding free energies using the molecular mechanics energies combined with the generalized Born (GB) and surface area continuum solvation (MM-GBSA) method [51]. The average of gas-phase and solvation free energy values were taken throughout 300 ns simulations. The calculations were performed by CHARMM36 programming program. The average binding free energy is formulated as a sum of the gas phase contribution, the solvation energy contribution and the entropic contribution which is shown as:

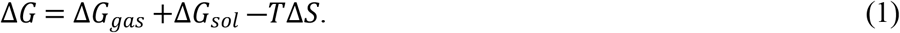

The change in binding energy was calculated with the following formula for K-Ras4B dimer systems:

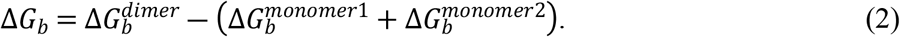

## Results

### Selection of the mutant K-Ras4B dimeric systems

In the initial structures of α- and β-homodimer models generated by PRISM, we extracted the interface residues (**Table 1**) and defined the most critical residues with their corresponding energy scores calculated by FoldX for both interfaces (**Table 2**). These results show that some residues are more critical in dimerization. I21, I24, Q25, H27, Y40, and R41 are found to be computational hot spots at the β-homodimer, and H94, R97, L133, S136, and Y137 are found to be computational hot spots at the α-homodimer interfaces according to HotRegion. Based on our previous studies [19, 20], we selected E98R, K101A/R102A, and K101D/R102E mutations on the allosteric lobe interface, and S39/Y40A, R41A/K42A, and R41E/K42D mutations on the effector lobe interface. In our previous studies [20], we tested these mutants *in vitro* using the Bimolecular fluorescence complementation (BiFC) system, investigating the Ras-Ras interactions in HEK293T cells. When there was an interaction between Ras proteins, a strong fluorescence signal was expected. The signals were mainly around the plasma membrane where K-Ras4B was located, suggesting the K-Ras4B proteins interact. The cells expressing K101D/R102E double mutants (on top of G12D) yielded less fluorescence signals compared to the cells expressing solely (K-Ras4B^G12D^, suggesting that K101/R102 residues play a role in interaction between K-Ras4B proteins [20]. According to our BiFC experiments, we selected two mutations with opposite charge, K101D/R102E for the allosteric lobe interface and R41E/K42D, for the effector lobe interface, and introduced them to the oncogenic KRas4B^G12D^ α- and β-homodimers, respectively. There were four simulation systems containing two allosteric lobe interface dimers, oncogenic K-Ras4B^G12D^ and mutant K-Ras4B^G12D/K101D/R102E^ α-homodimers, and two effector lobe interface dimers, oncogenic K-Ras4B^G12D^ and mutant K-Ras4B^G12D/R41E/K42D^ β-homodimers.

**Table 1.** Interface residues defined in the α- and β-homodimer structures.

| **K-Ras4B-GTP** | **Interface residues** |
| --- | --- |
| **α-homodimer** | E91, H94, H95, R97, E98, K101, R102, D105, S106, E107, K128, L133, R135, S136, Y137 |
| **β-homodimer** | I21, I24, Q25, H27, V29, E31, D33, I36, E37, D38, S39, Y40, R41, K42, Q43, L52 |

**Table 2.** K-Ras4B dimer binding free energy calculation by FoldX.

| K-Ras4B-GTP | Mutations | $\Delta\Delta G$ (kcal/mol) |
| --- | --- | --- |
| **β-homodimer** | S39A | 0.0756 |
|  | Y40A | 3.001 |
|  | S39A/Y40A | 3.069 |
|  | R41E | 9.931 |
|  | K42D | 5.288 |
|  | R41E/K42D | 15.894 |
|  | R41A | 3.251 |
|  | K42A | 3.192 |
| **α-homodimer** | R41A/K42A | 7.168 |
|  | E98R | 8.04 |
|  | E98Q | 3.393 |
|  | K101D | 7.22 |
|  | R102E | 5.587 |
|  | K101D/R102E | 12.105 |
|  | E98A | 3.831 |
|  | K101A | 4.266 |
|  | R102A | 3.844 |
|  | K101A/R102A | 6.49 |

### Stabilities of the oncogenic K-Ras4B^G12D^-GTP α- and β-homodimers

During the simulations, we observed that both oncogenic K-Ras4B^G12D^-GTP α- and β-homodimers are stable. As can be seen from the time-series of snapshots (**Fig 3**), no immediate dissociation of the dimers was monitored. However, we encountered large fluctuations in the Switch I and II regions during the simulations (**S1A Fig**). Of interest noticed for the oncogenic β-homodimer is that one of the K-Ras4B monomer yielded relatively larger fluctuations in the Switch I region than the other monomer. Large fluctuations of the Switch I loop are eminent when compared to the mutant β-homodimer (**S2 Fig**). The fluctuations induce conformational changes of the Switch I loop, which oscillates between the closed and open catalytic site conformations. Similar observations were reported for both wild-type and mutant H-Ras-GTP in the open and closed states using the MD simulations and crystallography experiments [52].

**Fig 3.**
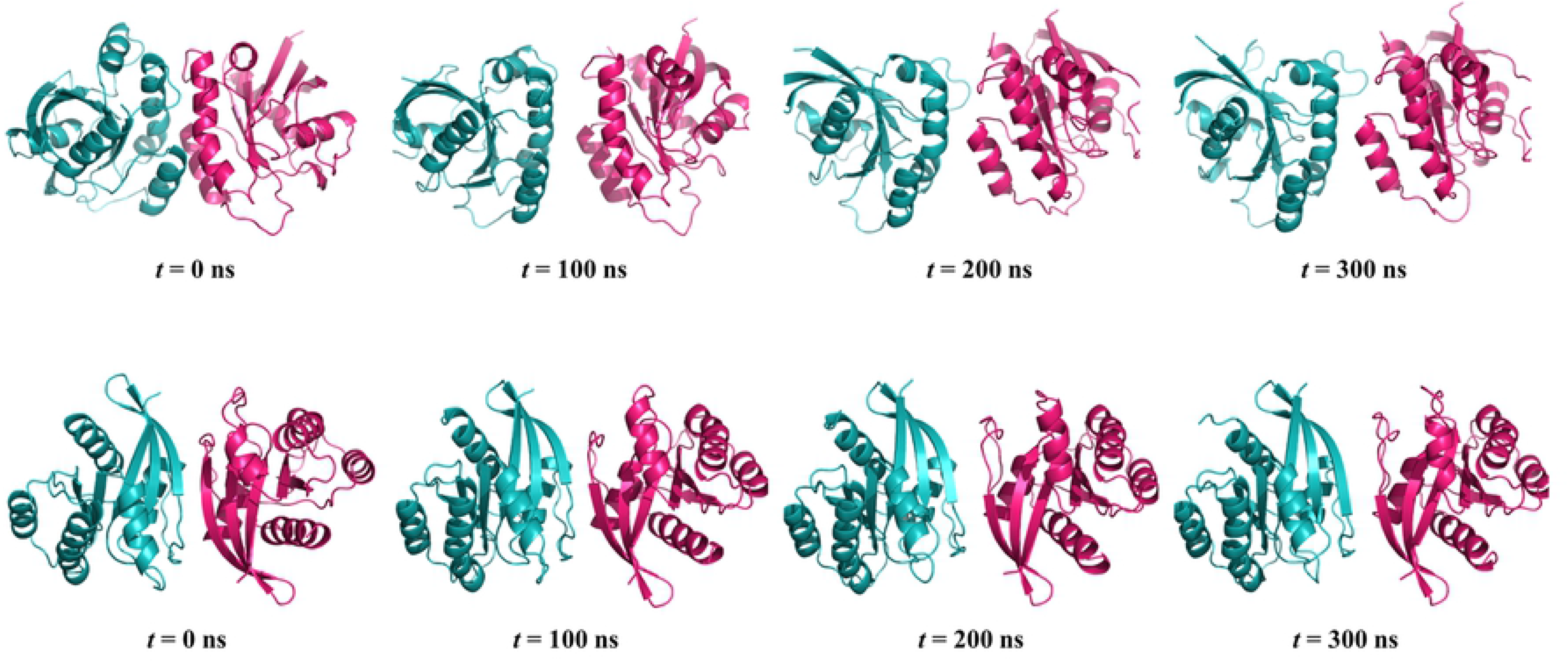
Simulated systems of the oncogenic K-Ras4B^G12D^ dimer complex. Time-series of the oncogenic α-homodimer (upper row), and that of the oncogenic β-homodimer.

For the oncogenic α-homodimer, the salt bridge interactions are a major driving force in stabilizing the dimer complex. Immediate drifting away of the proteins from the complex can be prevented due to strong salt-bridge interactions between the K-Ras4B monomers. To identify intermolecular interacting residue pairs that are responsible for the dimeric association, the atomistic interactions at the interfaces were investigated. We observed significant intermolecular salt bridge interactions at the interfaces (**Table 3**). For the oncogenic α-homodimer, E107-K101 and K128-E91 are the most frequently observed pairs of the salt bridge interaction at the interface. For the oncogenic β-homodimer, D37-K41 and D33-K42 are the most frequently observed pairs of the salt bridge interactions at the interface. In addition to the salt bridge formation, the intermolecular backbone hydrogen bonds (H-bonds) add to the stability of the β-homodimer. The H-bond interactions formed by the interacting pairs, S39-S39 and D37-K41, strongly retain the β-sheet dimer interface. These residues constitute the shifted β-sheet extension interface, consistent with previous observations [19, 21].

**Table 3.** Salt bridge formations throughout the simulations.

| Dimer systems | Salt bridge interacting pairs<br>M1 - M2 (%) |
| --- | --- |
| **K-Ras4B<sup>G12D</sup>**<br>**α-homodimer** | E107 - K101 (34.8)<br>K128 - E91 (38.3)<br>E98 - K101 (18.5)<br>K101 - E98 (6.3) |
| **K-Ras4B<sup>G12D</sup>**<br>**β-homodimer** | D38 - K42 (51.2)<br>K42 - D33 (47.5)<br>E37 - R41 (41.9)<br>D33 - K42 (40.9) |
| **K-Ras4B<sup>G12D/K101D/R102E</sup>**<br>**α-homodimer** | D126 - K128 (7.6)<br>K128 - D126 (6.6)<br>E91 - R135 (7.0)<br>R135 - E91(6.0) |

Salt bridge formations throughout the simulation trajectories for the oncogenic K-Ras4B<sup>G12D</sup> $\alpha$ - and β-homodimers, and for the mutant K-Ras4B<sup>G12D/K101D/R102E</sup> α-homodimer. M1 and M2 denotes monomer 1 and 2, respectively.

### Comparisons of oncogenic K-Ras4B^G12D^ dimers with mutant K-Ras4B^G12D/K101D/R102E^ and K-Ras4B^G12D/R41E/K42D^ dimers

To investigate stabilities of the K-Ras4B dimeric systems, we calculated the center of mass distance between two monomers in each dimer (**Fig 4**). For the oncogenic α-homodimer, the center of mass distance is measured ~35 Å, although large fluctuations in the distance are observed at *t* ~ 130 ns. The fluctuations occur due to rearrangement of the helices at the dimer interface, resulting that the dimer slightly alters the interface, shifting to an asymmetric helical interface (**Fig. 3**). In contrast, the oncogenic β-homodimer stably maintains the center of mass distance ~33 Å throughout the simulation. For the mutant α-homodimer, we also observed large fluctuations in the distance at *t* ~ 230 ns due to rearrangement of the allosteric helices. Similar to the oncogenic α-homodimer, the mutant α-homodimer also yields the asymmetric helical interface, retaining the dimeric association. However, for the mutant β-homodimer, we observed that the dimer is not stable, separated into two monomers at the early stage of the simulations. The separation is caused by the electrostatic repulsions between β2 strands, exerted from the mutated residues with opposite charges. After the separation, each separated monomer is stable, exhibiting less fluctuations in the Switch I and II regions as compared to those in the dimeric complex (**S1 Fig**).

**Fig 4.**
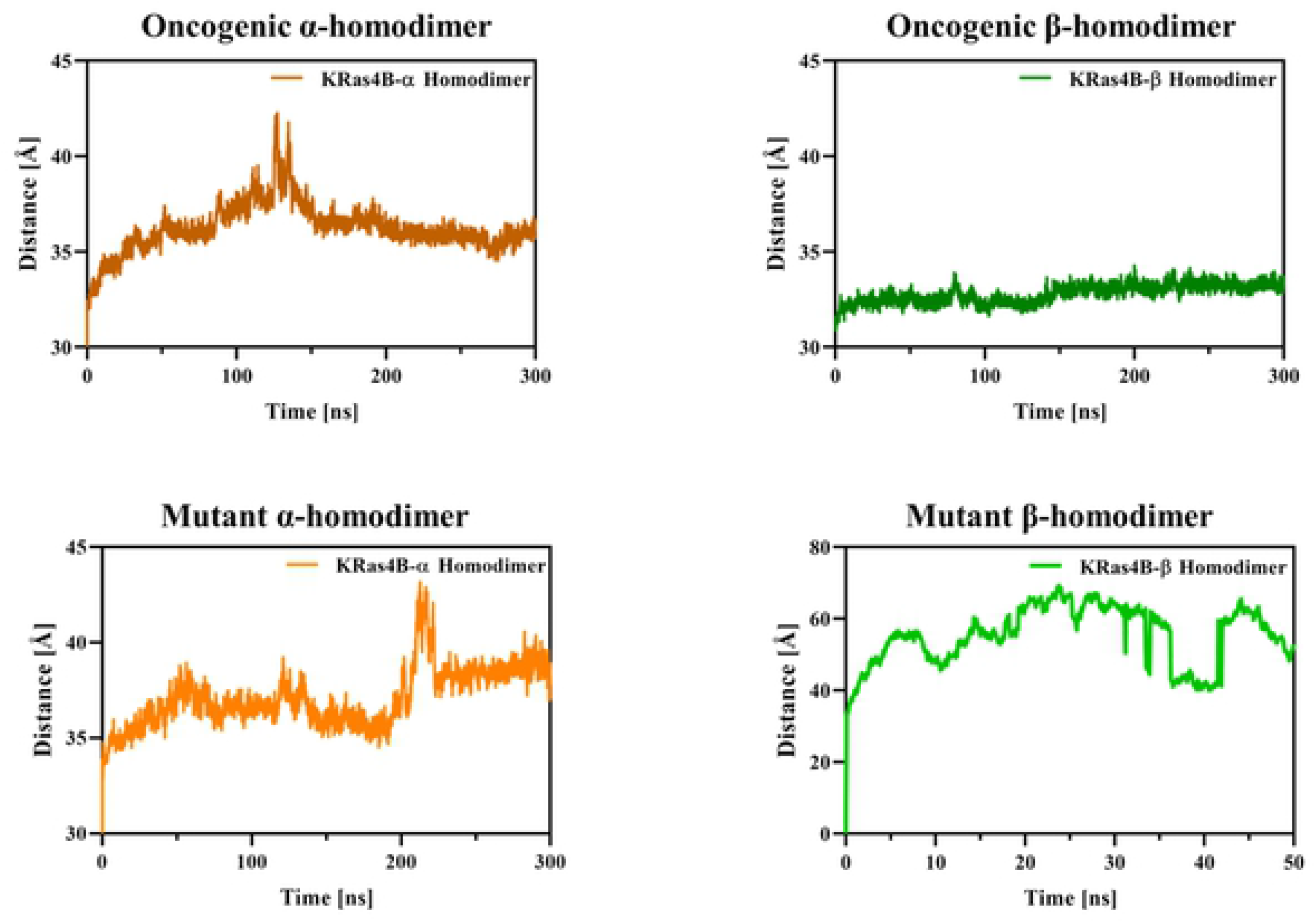
The center of mass distance. Time series of the center of mass distance between two monomers for the oncogenic K-Ras4B^G12D^ α- and β-homodimers (upper panels), and that for the mutant K-Ras4B^G12D/K101D/R102E^ α-homodimer and K-Ras4B^G12D/R41E/K42D^ β-homodimer (lower panels).

To identify intermolecular interacting residue pairs for the mutant α-homodimer, we also examined the atomistic interactions at the interfaces (**Table 3**). We only investigated the residue pairs for the mutant α-homodimer, since the mutant β-homodimer is separated, thus no interface residue pairs. The residue pairs for the salt bridge interactions are different compared to those in the oncogenic α-homodimer. This suggests that helices are aligned in a different way at the interface, although both α-homodimers favor the similar asymmetric helical interface. We found that 6 % of the residues are observed to be conserved 90% of the time, 13% of the residues are observed to be conserved 70% of the time for the mutant α-homodimer.

### Clustering analysis for K-Ras4B homodimers

To provide the best representative model of K-Ras4B dimer complexes, we clustered the ensembles of the conformations over the simulation trajectories (**Fig 5**). We obtained 5 representative clusters for the oncogenic α-homodimer. The first and second clusters with 34.0% and 14.0% populations, respectively, exhibit the similar dimeric interactions using the α3-α4-α5 helices from one monomer and the α2-α3 helices from the other monomer. Unlike the conformations from two highly populated clusters, the representative conformations from next three less populated clusters are similar to each other. The initial dimeric interface is formed by the symmetric α3-α4/α3-α4 helical alignment. During the course of the simulation, the symmetric helical alignment (with 31% population) is steadily converted to an asymmetric α3-α4-α5/α2-α3 helical alignment (with 48% population) (**S3 Fig**). The dimer adopts the asymmetric helical alignment using the α3-α4/α3 and α5/α2 interfaces. The asymmetric α3-α4/α3 helical interface is commonly observed in the K-Ras4B dimer with the allosteric lobe interface [21, 22]. The occurrence frequency of residue pairs that contributes interface formation in the oncogenic α-homodimer are given in **S1 Table**. We also obtained 5 representative clusters for the oncogenic β-homodimer, and found that unlike the α-homodimer, the representative conformations from each cluster are similar to each other. The oncogenic β-homodimer retains the shifted β-sheet extension interface with relatively high affinity. We summarized the occurrence frequency of residue pairs that contributes interface formation in the oncogenic β-homodimer in **S2 Table**.

**Fig 5.**
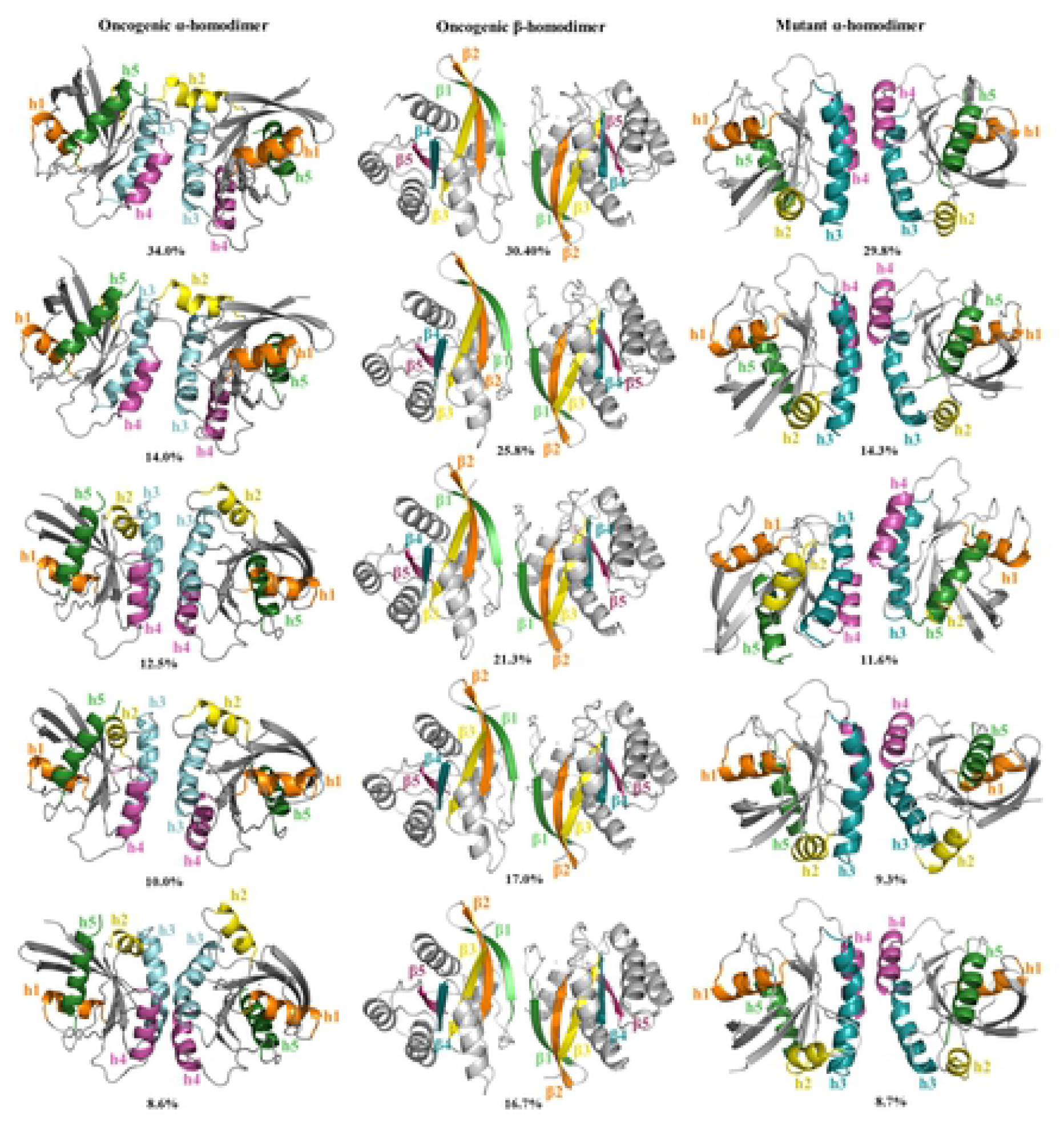
Clustering analysis. Snapshots and populations of the five representatives for the most populated conformational clusters for the oncogenic K-Ras4B^G12D^ α-homodimer (left column) and β-homodimer (middle column). The same for the mutant K-Ras4B^G12D/K101D/R102E^ α-homodimer (right column).

For the mutant α-homodimer, we provide 5 clusters representing the best models for the mutant dimeric complex (**Fig 5**). No clusters for the mutant β-homodimer were obtained due to separation. Similar to the oncogenic α-homodimer, the mutant α-homodimer also adopts the asymmetric helical alignment using the α3-α4/α3 interface, abandoning the initial symmetric α3-α4/α3-α4 interface (**S4 Fig**). However, no α5/α2 interface was observed. The representative clusters were sampled during the simulation in the order of their emergences, cluster 4 → cluster 5 → cluster 1 → cluster 2 → cluster 3.

### Binding energies for oncogenic K-Ras4B^G12D^ dimers

The binding free energies of the dimer systems were calculated by using the MM-GBSA method. The binding free energy of the oncogenic α-homodimer seems to be less favorable than the oncogenic β-homodimer (**Table 4**), suggesting that the allosteric lobe interface is not strong as the effector lobe interface, consistent with the wild-type case [19, 21]. The α-homodimer undergoes rearrangements of the allosteric helical interactions during the simulations, contributing to the enthalpy changes for individual clusters. We observed that, indeed each cluster has considerably different enthalpy changes for the oncogenic α-homodimer. These are specifically −54.8, −50.9, −44.1, −40.2 and −65.3 kcal/mol for cluster 1-5, respectively. Therefore, the initial complex was the most favorable one, then the energy increased and after 150 ns, it became stable for clusters 1 and 2 (**S5 Fig**). In contrast, the enthalpy change is relatively stable for the oncogenic β-homodimer.

**Table 4.** MM-GBSA results for the oncogenic K-Ras4B^G12D^ dimer complexes.

| K-Ras4B<sup>G12D</sup> | $\Delta H$ (kcal/mol) | $-T\Delta S$ (kcal/mol) | $\Delta G$ (kcal/mol) |
| --- | --- | --- | --- |
| α-homodimer | $-49.6 \pm 13.7$ | 71.0 | 22.1 |
| β-homodimer | $-68.1 \pm 8.2$ | 75.9 | 7.8 |

### Possible higher order complexes for the alpha control

K-Ras4B forms a functional dimer through the allosteric lobe interface. The α3-α4/α3-α4 helical alignment was defined as a major K-Ras4B allosteric lobe interface, while the α4-α5/α4-α5 helical alignment was appeared to be a minor interface [19, 21, 22, 53]. We provide some possible higher order homo-complexes for the oncogenic α-homodimers (**Fig 6, S6 Fig**). We used the cluster representatives from the **Fig 5** (left column) as well as the minor α-homodimer structure using the α4-α5/α4-α5 helical interface from previous studies [19]. We construct some possible trimers, tetramers, and pentamers, which are obtained by the superimposition of the dimers. We selected the complex structures that have the C-termini facing to the same surface, where each monomer will be bound to the membrane. All homo-complexes were generated by HSYMDOCK web-server [54]. As an initial input, we used representative 1 and representative 3, 47.6% of all visited conformations. Representative 2 is very similar to representative 1 (iRMSD < 1.614 Å); therefore, we did not consider it. Interface residues are obtained by HotRegion web-server [41], and corresponding helices were indicated. According to our results, trimer, tetramer, and pentamer formation occur via α3-α4 and α5 interfaces, exposing the effector binding sites. When we tried to form the higher order oligomers starting with the cluster representatives of the mutant α-homodimers, we could not manage to obtain regular trimers, tetramers or pentamers. This might suggest that although the mutant α-homodimer is plausible, it is not possible to construct higher order structures from them.

**Fig 6.**
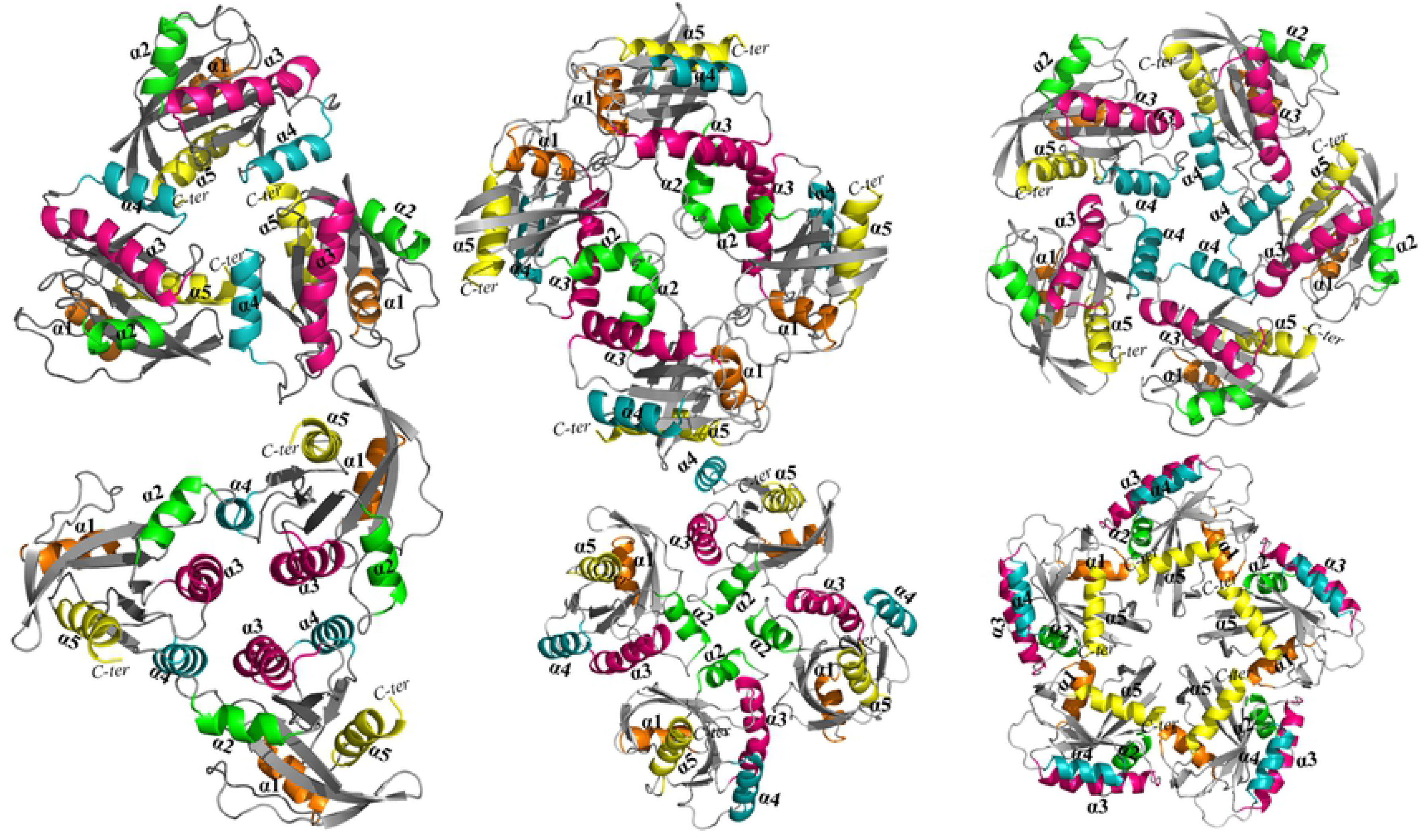
The predicted higher order homo-complexes. The trimer, tetramer and pentamer complexes using symmetry operations with the oncogenic K-Ras4B^G12D^ α-homodimer.

## Discussion

Monomeric Ras can bind Raf, however Raf should act as a dimer, which can be facilitated by Ras dimerization [18, 21]. Using *in silico* and *in vitro* methods, we previously demonstrated that wild-type K-Ras4B in the GTP-bound state can dimerize through two major interfaces involving the allosteric and effector lobe interfaces [19, 21, 22]. The allosteric lobe interface yields a functional α-homodimer, since the effectors can bind to the exposed effector binding site. In contrast, the effector lobe interface produces a nonfunctional β-homodimer, since the dimer shares the same interface with the effectors. K-Ras4B favors to form a major α-homodimer using α3 and α4 helices at the allosteric lobe, but the population of a dimer involving α4 and α5 helices is low. A major K-Ras4B β-homodimer contains a shifted β-sheet extension between β2 strands. The β-homodimer exhibits relatively higher affinity than the α-homodimer. A minor K-Ras4B β-homodimer reveals a β-sandwich interface involving side-chain interactions of β1, β2, and β3 strands. The β-sandwich interface emerged from the exact β-sheet alignment due to H-bonds mismatch between β2 strands [19, 21]. A similar nonfunctional β-sandwich dimer stabilized by two BI-2852 molecules was recently discovered [35–37].

Our oncogenic K-Ras4B^G12D^ α-homodimer retains the dimeric association. The oncogenic α-homodimer favors an asymmetric helical alignment using the α3-α4/α3 and α5/α2 interfaces, abandoning the symmetric α3-α4/α3-α4 helical alignment. We observed that the asymmetric α3-α4/α3 helical interface is popular among K-Ras4B dimers with the allosteric lobe interface [21, 22]. The oncogenic β-homodimer is relatively stable during the simulations, preserving the same interface as observed in the wild-type simulations [19, 21]. The mutant Ras4B^G12D/K101D/R102E^ α-homodimer marginally preserves the allosteric lobe interface using the similar asymmetric α3-α4/α3 helical interface as observed for the oncogenic case, while the mutant K-Ras4B^G12D/R41E/K42D^ β-homodimer is dissociated at the early in the simulations. The asymmetry in the conformation of functional Ras dimer may help to deduce the shape of the nanoclusters, the higher order homo-complexes, suggesting that it is less likely linear but more likely curved or circular. Using the HSYMDOCK web-server [54], we delineate K-Ras4B nanoclusters as the trimeric, tetrameric, pentameric shapes using the oncogenic K-Ras4B^G12D^ α-homodimer interface. However, no higher order homo-complex is predicted for the mutant Ras4B^G12D/K101D/R102E^ α-homodimer due to the weak dimeric interface.

The β-homodimer with the effector lobe interface overlaps with the binding region of its effectors, whereas the α-homodimer with the allosteric lobe helical interface is believed to promote Ras dimerization [20, 22] and thus Raf dimerization. Recent site-directed mutagenesis and cellular localization experiments showed that K101D/R102E double mutations on the allosteric lobe of K-Ras4B^G12D^ reduce dimerization at the plasma membrane and slightly decrease downstream phosphorylated extracellular signal-regulated kinase (ERK) levels [20]. In contrast, R41E/K42D double mutations on the effector lobe retain dimerization at the plasma membrane but completely abrogate ERK phosphorylation. Both double mutations increase Akt phosphorylation. The altered phosphorylation levels on the downstream effectors are composite results of the mutations affecting the Ras dimerization and the HVR dynamics interrupting the Ras interaction at the plasma membrane. In line with the experiments, the mutant K-Ras4B^G12D/K101D/R102E^ α-homodimer exhibits relatively weak dimer interface in solution, and thus reducing dimerization and decreasing pERK levels at the plasma membrane. On the other hand, the mutant K-Ras4B^G12D/R41E/K42D^ β-homodimer is unstable in solution, but the experiments verified dimerization at the plasma membrane. This indicates that the mutant avoids unfavorable effector lobe interface, instead promoting dimerization through the allosteric lobe interface at the membrane. However, it was observed that the mutant K-Ras4B^G12D/R41E/K42D^ mutant blocks ERK phosphorylation in the MAPK pathway, since it cannot activate Raf-1 [20].

Our simulations verified that the major effector lobe interface is made of a single state and the major allosteric lobe interface has several states. Although the effector lobe interface is more stable than the allosteric lobe interface, the functional dimeric interface is through the allosteric lobe interface containing α3 and α4 helices with the exposed effector binding sites for recruiting two Rafs to the plasma membrane [21, 22]. The multiple interfaces observed for the allosteric lobe interface might help to draw functional K-Ras4B nanoclusters.

## Acknowledgements

This work has been supported by TUBITAK Research Grant No: 114M196. This project has been funded in whole or in part with federal funds from the National Cancer Institute, National Institutes of Health, under contract HHSN26120080001E. The content of this publication does not necessarily reflect the views or policies of the Department of Health and Human Services, nor does mention of trade names, commercial products, or organizations imply endorsement by the U.S. Government. This Research was supported [in part] by the Intramural Research Program of the NIH, National Cancer Institute, Center for Cancer Research. Computations have been performed at the high-performance center, Koc University and at the high-performance computational facilities of the Biowulf PC/Linux cluster at the National Institutes of Health, Bethesda, MD (https://hpc.nih.gov/).

## Author Contributions

K.K. and M.E.O. performed the molecular dynamics studies. K.K. analysed data, prepared tables and figures. H.J. guided the simulations. R.N., H.J. A.G. and O.K. designed and conceptualized the project. H.J., K.K., M.E.O., O.K. and A.G. performed the article writing. H.J, and O.K. made critical revisions and approved final version. All of the authors reviewed and approved the final manuscript.

**Supporting information**

**S1 Fig. The root-mean-square fluctuation (RMSF) of each system**. (A) The RMSF of the oncogenic K-Ras4B^G12D^ α- and β-homodimers, and (B) the same of the mutant K-Ras4B^G12D/K101D/R102E^ α-homodimer and K-Ras4B^G12D/R41E/K42D^ β-homodimer.

**S2 Fig. Fluctuations in the Switch I regions**. Switch I regions of the oncogenic K-Ras4B^G12D^ β-homodimer (left panel) as compared to that of the mutant K-Ras4B^G12D/R41E/K42D^ β-homodimer.

**S3 Fig. Highly populated clusters representing the conversion of interfaces**. Interfaces shifted from the symmetric α3-α4/α3-α4 helical alignment (with 31% population) towards the asymmetric helical alignment involving the α3-α4/α3 and α5/α2 interfaces (with 48% population) for the oncogenic K-Ras4B^G12D^ α-homodimer.

**S4 Fig. Snapshots representing the conversion of interfaces**. Snapshots representing the conversion of interface from the symmetric α3-α4/α3-α4 helical alignment to the asymmetric α3-α4/α3 helical alignment for the mutant K-Ras4B^G12D/K101D/R102E^ α-homodimer (upper panels), and the dissociation of the mutant K-Ras4B^G12D/R41E/K42D^ β-homodimer (lower panels).

**S5 Fig. Time series of the enthalpy changes, Δ*H*, during the simulations for the oncogenic K-Ras4B^G12D^ α-homodimer (upper panel) and β-homodimer (lower panel).**

**S6 Fig. Some examples of the predicted higher order homo-complexes**. The trimer, tetramer and pentamer complexes using symmetry operations with the oncogenic K-Ras4B^G12D^ α-homodimer and the interface residues together with the secondary structure elements in these complexes are listed.

